# A spatiotemporal steroidogenic regulatory network in human fetal adrenal glands and gonads

**DOI:** 10.1101/2021.12.22.473776

**Authors:** Yifu Wang, Bingqian Guo, Yajie Guo, Nana Qi, Yufang Lv, Yu Ye, Yan Huang, Xinyang Long, Hongfei Chen, Cheng Su, Liying Zhang, Qingyun Zhang, Minxi Li, Jingling Liao, Yunkun Yan, Xingning Mao, Yanyu Zeng, Jinghang Jiang, Zhongyuan Chen, Yi Guo, Shuai Gao, Jiwen Cheng, Yonghua Jiang, Zengnan Mo

## Abstract

Human fetal adrenal glands produce substantial amounts of dehydroepiandrosterone (DHEA), which is one of the most important precursors of sex hormones. However, the underlying biological mechanism remains largely unknown. Herein, we sequenced human fetal adrenal glands and gonads from 7 to 14 GW via the 10× Genomics single-cell transcriptome techniques and reconstructed their location information by Spatial Transcriptome, conducted COOL-seq for the *MC2R+* inner zone steroidogenic cells during the time window of sex differentiation (8-12GW). We found that relative to gonads, adrenal glands begin to synthesize steroids early. The coordination among steroidogenic cells and multiple nonsteroidogenic cells promotes adrenal cortex construction and steroid synthesis. Notably, during the time window of sex differentiation (8–12 GW), key enzyme gene expression shifts to accelerate DHEA synthesis in males and cortisol synthesis in females. Furthermore, high *SST*+ expressions in the adrenal gland and testis amplify androgen synthesis in males. Our research highlights the robustness of the action of fetal adrenal glands on gonads to modify the process of sexual differentiation.

## Introduction

Human fetal adrenal glands are huge endocrine organs composed of a definitive zone (DZ), a fetal zone (FZ) (approximately 75%), and a transition zone (TZ) (after 14 gestational weeks [GW]) that produce huge amounts of dehydroepiandrosterone (DHEA) and cortisol(Ishimoto & Jaffe, 2011, Melau, Nielsen et al., 2019). DHEA is the most prominent precursor for sex hormone(Melau et al., 2019).. DHEA is clearly the main source of placenta estradiol (E2), which plays a critical role in regulating the development and maturation of fetal and maternal–fetal interface(Ishimoto & Jaffe, 2011, Miller & Auchus, 2019). In testis or ovary, DHEA is processed to become a different subtype of testosterone or estrogen, which drives sexual differentiation in organs(Asby, Arlt et al., 2009, Ishimoto & Jaffe, 2011, Melau et al., 2019).

The human fetal gonads are “bipotential” before 8 GW(Rotgers, Jorgensen et al., 2018).. Gonads regulate the sexual differentiation of external reproductive organs by synthesizing sex hormones in a dose-dependent manner(Christou & Tigas, 2018, Rotgers et al., 2018, Scott, Mason et al., 2009). Extensive evidence shows that the dysfunction of fetal adrenal glands leads to reproduction-related diseases, such as polycystic ovary syndrome(Goodarzi, Carmina et al., 2015), impaired spermatogenesis, and testicular atrophy(Scott et al., 2009). If the malfunction in adrenal glands is caused by mutations in various genes, such as *CYP21A2*, *CYP11B1*, and *POR*, and when adrenal dysfunction occurs before the window of sexual differentiation, the malfunction may cause serious virilization or feminization(Asby et al., 2009, Miller & Auchus, 2011). On account of DHEA circulation in fetal adrenal glands and gonads, we hypothesized that adrenal glands and the steroidogenesis regulatory network between adrenal glands and gonads may play a critical role in initial sexual development during the time window of sex differentiation.

However, investigating the functions of human fetal adrenal glands is difficult for two reasons. First, the origin of circulating DHEA and adrenal-derived androgens in humans and nonhuman primates is distinct from that in other mammalian species(Abbott & Bird, 2009, Ishimoto & Jaffe, 2011). Second, adrenal glands are a complex structure with various cellular component organs consisting of numerous heterogeneous cell types; most cortical cells originate from the mesoderm, neurocytes from the ectoderm(Dong, Yang et al., 2020, Furlan, Dyachuk et al., 2017), and germ cells from the yolk sac(Upadhyay & Zamboni, 1982). The precise composition of tissue structures and functions of fetal adrenal glandsare unclear.

The recent development of single-cell sequencing technology has allowed the determination of complicated landscape of cell biology in the tissue(Bian, Gong et al., 2020, Dong et al., 2020, Furlan et al., 2017, Li, Dong et al., 2017, Tang, Barbacioru et al., 2009). Through the joint use of sc-RNA sequencing, Spatial Transcriptomics and multi-omics sequencing technology (COOL-seq)(Guo, Li et al., 2017, Satija, Farrell et al., 2015, Stahl, Salmen et al., 2016), we can make a comprehensive and thorough analysis of the spatial and temporal information within the organization from the dimension of multi-omics. In this study, we draw a fine map of the adrenal glands and gonads of 7-14 GW human fetal, which provide an unprecedented resource to understand the complexity of adrenal glands and shed light on the action of fetal adrenal glands and gonads to modify the process of sexual differentiation.

## Results

### Expression Programs of Cell Lineages in Fetal Adrenal Glands

To generate a comprehensive landscape of steroid hormones in fetal adrenal glands and gonads spanning the window of sexual differentiation, adrenal glands and gonads (7–14 GW) samples were collected. The samples were digested and subjected to single-cell transcriptome sequencing by using the 10× Genomics system. Frozen sections from 8-9 GW fetal adrenal tissues were used for spatial transcriptome. The samples were classified into three stages: before (<8 GW), within (8–12 GW), and after the window of sexual differentiation (>12 GW)(Hanley & Arlt, 2006) (Fig 1A). A total of 75,482 adrenal cells and 53,508 gonad cells passed the standard quality control. Filtered out erythrocytes were retained for downstream analysis (Table EV1).

**Figure 1.**
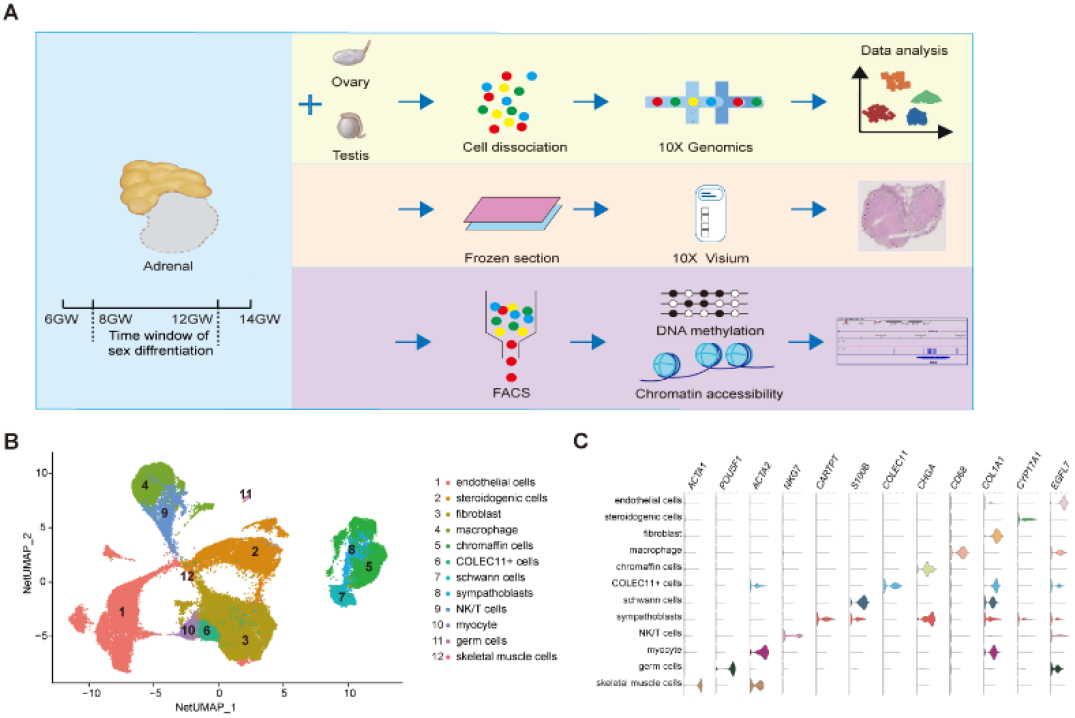
Global patterns of single-cell expression profiles of fetal adrenal glands. A. Experimental schematic; 10 adrenal glands and 8 gonads in 10× Genomics; 2 adrenal glands in Spatial transcriptome, 10 adrenal glands in COOL-seq. B. Uniform manifold approximation and projection analysis (UMAP) of the transcriptomes of fetal adrenal cells in 10× Genomics data (n = 75,482). The clusters were identified by marker genes, as shown in C and D. C. Violin plot overview of the expression of selected marker genes by the fetal adrenal clusters. Detailed cell information and differentially expressed genes can be found in Table EV2.

On the basis of 10× Genomics data, uniform manifold approximation and projection analysis (UMAP) revealed 12 main cluster groups (Fig 1B and C), including the cells that mainly perform steroidogenic functions that previous studies defined as steroidogenic cells (CYP17A1+)(Ishimoto & Jaffe, 2011), as well as other cells, such as neurocytes (CHGA+ chromaffin cells, CARTPT+ sympathoblasts, S100B+ Schwann cells)(Furlan et al., 2017) and immune cells (CD68+ macrophage, NKG7+ NK/T cells)(Bian et al., 2020) (Fig 1C; Table EV2).

### Spatiotemporal transcriptome characteristics of steroidogenic cells

Steroidogenic cells, classically defined by *CYP17A1*, are a manifestation of adrenal function for steroid biosynthesis(Miller & Auchus, 2011). We performed an unsupervised analysis of the gene expression profiles of these cells. Eight subtypes of steroidogenic cells were identified (Fig 2A and B; Table EV3). The gene expression heat map showed that the relative expression levels of the genes known to be associated with cyst stem cells (*VSNL1* and *NOV* in T3 and *COL1A1* and *RSPO3* in T4), proliferation (*MKI67*, *TOP2A* in T2), and steroidogenic metabolism genes (*CYP11B1* and *SULT2A1* mainly in T3 and T5) varied in the eight subtypes (Fig 2B). T2 had a high cell cycle score defined as proliferative cells (Fig EV1A). Gene ontology (GO) analysis revealed that chromosome segregation, organelle fission, and nuclear division were enriched in subtype T2 (Fig EV1C). *NOV* is a marker gene for DZ, it is highly expressed in T3; *SULT2A1* and *CYP11B1* were highly expressed in T1 and T5 which characterized as FZ (Fig 2B; Table EV3). Immunostaining showed that cells expressing *NOV*, *MC2R* and *CYP17A1* were sequentially distributed from DZ to FZ (Fig EV1B). The characteristics of T4 cells were similar to those of fibroblasts (Fig 2B; Table EV3) and the steroidogenic genes were limited, GO analysis revealed that extracellular matrix organization, extracellular structure organization, and axonogenesis were enriched in this subtype (Fig EV1C), indicating that T4 may be involved in forming the adrenal zone.

**Figure 2.**
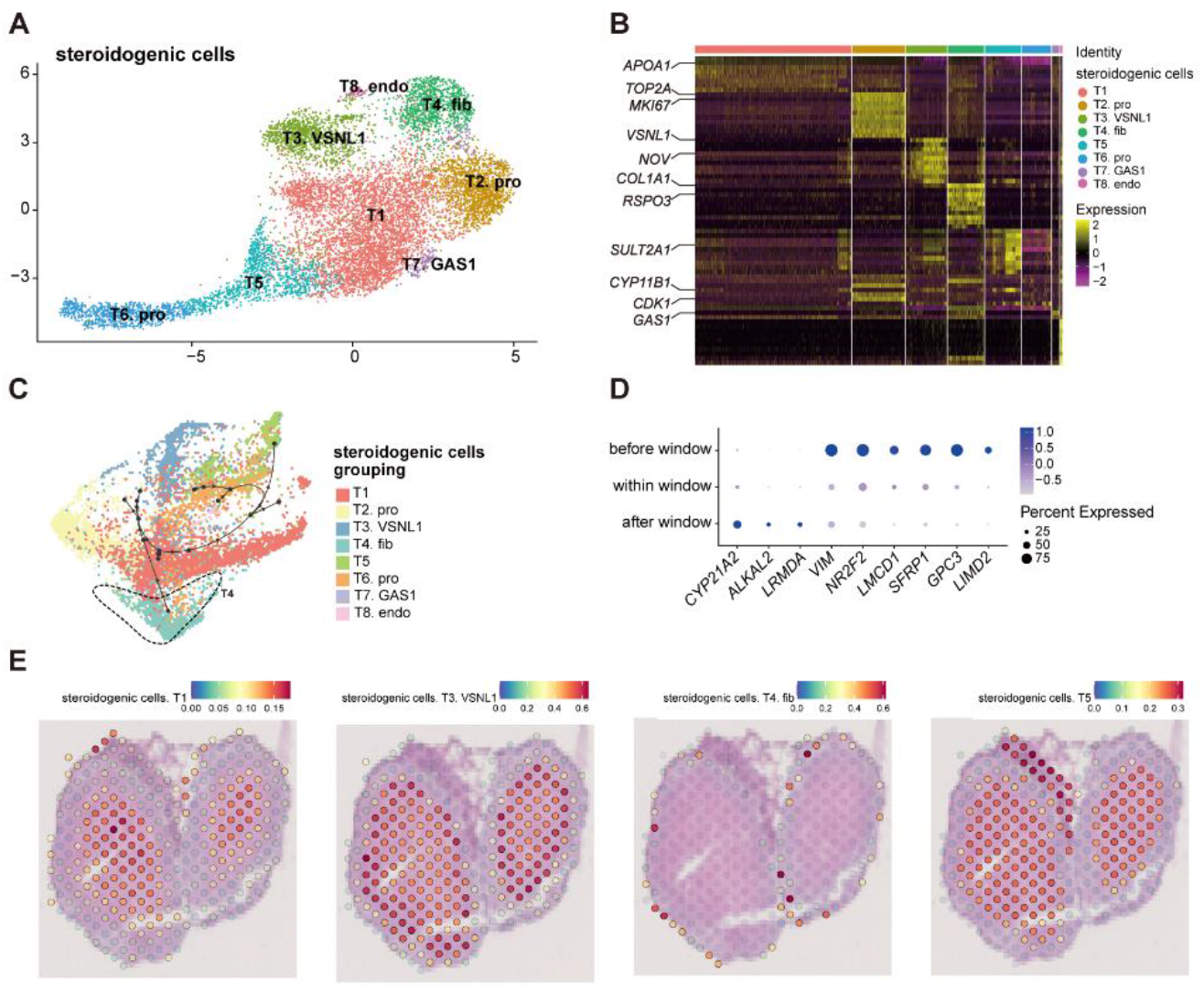
Dynamic gene expression patterns of fetal adrenal gland steroidogenic cells. A. Uniform manifold approximation and projection analysis visualization of professional steroidogenic cells for 10× Genomics data (n = 10,651). B. Heat map of the top 10 differentially expressed genes (DEGs) between professional steroidogenic cell populations (n = 10,651). Color scale: yellow, high expression; purple, low expression. Detailed cell information and DEGs can be found in Table EV3. C. Differentiation of trajectory of professional steroidogenic cells using Dyno. Arrow direction indicates the trajectory of cell differentiation. D. Dot plots of differential gene expression of three-stage professional steroidogenic cell groups (before, within, and after sexual differentiation). E. Visualization of the spatial transcriptome shows the location of T1, T3. *VSNL1*, T4. fib and T5 steroidogenic cells in 8GW fetal adrenals.

Spatial Transcriptome was used to reproduce the spatial positions of these types of steroidogenic cell(Stahl et al., 2016). The T1 and T5 steroidogenic cells were located in the FZ of the fetal adrenal; the T3 steroidogenic cells were mostly distributed in the outside of T1 and T5 steroidogenic cells, considered as the cells located in DZ. T4 steroidogenic cells, the progenitor cell as we defined, were only distributed in the peripheral location, which is under the cysts (Fig 2E and EV1D). In accordance with the classical subcapsular DZ to FZ centripetal differentiation model(Ishimoto & Jaffe, 2011), pseudotime analysis revealed a developmental trajectory of steroidogenic cells from T4 to other subtypes (Fig 2C). With the differentiation and maturity of steroidogenic cells, the steroidogenic-related genes *CYP17A1, CYP11B1* and *CYP21A2* were highly expressed (Fig 2C and EV1E). In summary, the single cell RNA sequencing data combined with Spatial Transcriptome enables us to support the classical model of the central differentiation and maturation of fetal adrenal steroidogenic cells from under the cyst.

The dot plots showed that along with fetal development, the motility regulator *LIMD2*, the lipid transport mediators *NR2F2* and *VIM*, and pluripotent genes (e.g., *GPC3*, *SFRP1*, and *LMCD1*) were progressively downregulated. Moreover, steroidogenic regulation-related genes, such as *LRMDA*, *ALKAL2*, and *CYP21A2*, were upregulated (Fig 2D). Steroidogenic cells with high expression of *CYP17A1* and *CYP11B1* were enriched in the innermost FZ while the cells high expresses *CYP21A2* were highly outside of them (Fig EV1F). Cells high expressing *HSD3B2* and *CYP11B2* were in the outermost part of the fetal adrenal glands (Fig EV1F). The *MC2R*, encoding the protein of ACTH receptor, is widely expressed in the FZ and TZ (Fig EV1F). Subsequently, *MC2R* was used as marker to sort the steroidogenic cells in inner zone.

### Steroidogenic Cells in Adrenal Glands Exhibit Distinct Expression Profiles across Different Sexes

Same as previously reported(Goto, Piper Hanley et al., 2006, Melau et al., 2019), a crest of *HSD3B2* within the sexual differentiation window only in females (Fig 3A and B). The temporarily elevated *HSD3B2* was thought to increase cortisone production and suppress DHEA synthesis through the competitive consumption of common precursors(Goto et al., 2006, Melau et al., 2019). However, the expression of another key enzyme for producing cortisol, *CYP21A2*, was limited in this period (Fig 2D), while *C*YP11B1 were detected. CYP11B*1* can not only produce cortisol, but also play a key role in the biosynthesis of 11-Oxygenated androgens, which are new highly bioactive androgen mainly synthesis in androgen (Davio, Woolcock et al., 2020, Pretorius, Arlt et al., 2017, Turcu, Rege et al., 2020).

**Figure 3.**
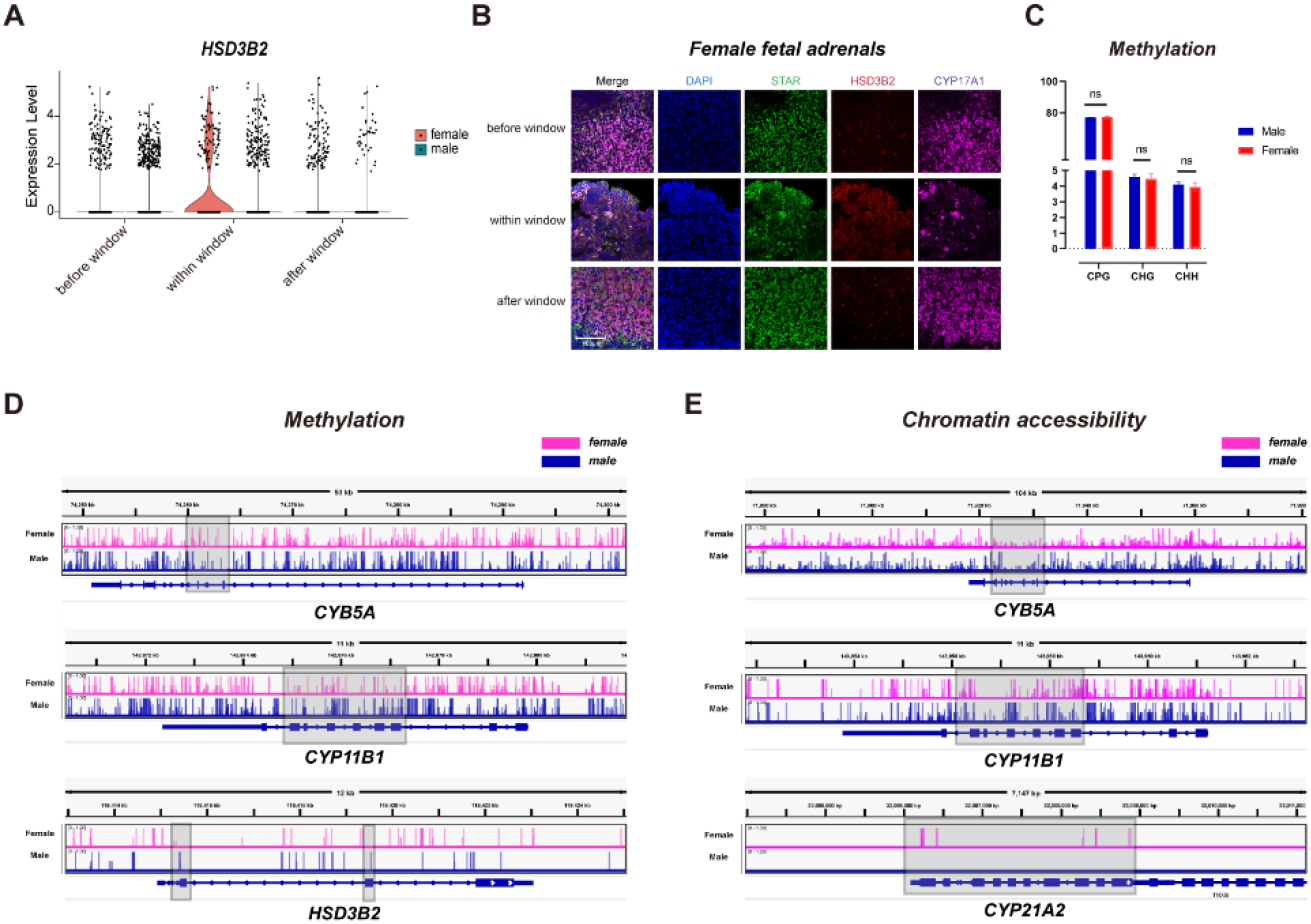
Dynamic gene expression patterns of fetal adrenal gland steroidogenic cells. A. Violin plots of *HSD3B2* gene expression patterns of female and male fetuses within the window of sexual differentiation. B. Immunofluorescence staining of StAR (green), HSD3B2 (red), and CYP17A1 (purple) in the female fetal adrenal glands spanning the window of sexual differentiation. Scale bar, 20 μm. n = 3. C. The overall methylation levels between male and female in COOL-seq. D. The landscape of methylation site in *CYB5A*, *HSD3B2*, *CYP11B1*, *CYP17A1* of human fetal adrenal *MC2R*+ steroidogenic cells (female, purple; male, bule). E. The landscape of chromatin accessibility site in *CYB5A*, *CYP11B1*, *CYP17A1*, *CYP21A2* of human fetal adrenal *MC2R*+ steroidogenic cells (female, purple; male, bule).

In order to explore whether sexual dimorphism exist in the epigenetic regulation of steroidogenic cells during the window of sexual differentiation, we conducted COOL-seq(Guo et al., 2017) on the adrenal *MC2R*+ steroidogenic cells (Fig 1A). The DNA methylation and chromatin state in those steroidogenic cells were detected. Although there were no significant differences in overall methylation levels between male and female, we found that the epigenetic profiles of the enzyme genes related steroid metabolizing are different. The methylation levels of exons in *HSD3B2* and *CYP11B1* were higher in males, while *CYB5A* was higher in females (Fig 3D). The chromatin accessibility of promoter regions is known to be highly associated with expression levels of the corresponding genes(Guo et al., 2017). *CYP21A2*, a key enzyme for producing cortisol was only can be accessed in female. Meanwhile, the *CYB5A* in male has more chromatin opening (Fig 3E). These results suggest that the steroidogenic cells in female are more likely to synthesize cortisol, on the other hand, most willing to synthesize DHEA in male.

Taken together, the data indicated the dynamic changes in the steroid synthesis function of the steroidogenic cells and revealed the sex-specific guiding mechanism in the window of sexual differentiation.

### Neurocytes in Fetal Adrenal Glands Participate in Steroid Regulation

To explore whether the fetal adrenal neurocytes have steroidogenic functions, the neurocyte group was further divided into 15 distinct subtypes, including 4 chromaffin cell subtypes, 2 sympathoblast subtypes, and 3 Schwann cell precursor (SCP) subtypes (Fig 4A and B; Table EV4).

**Figure 4.**
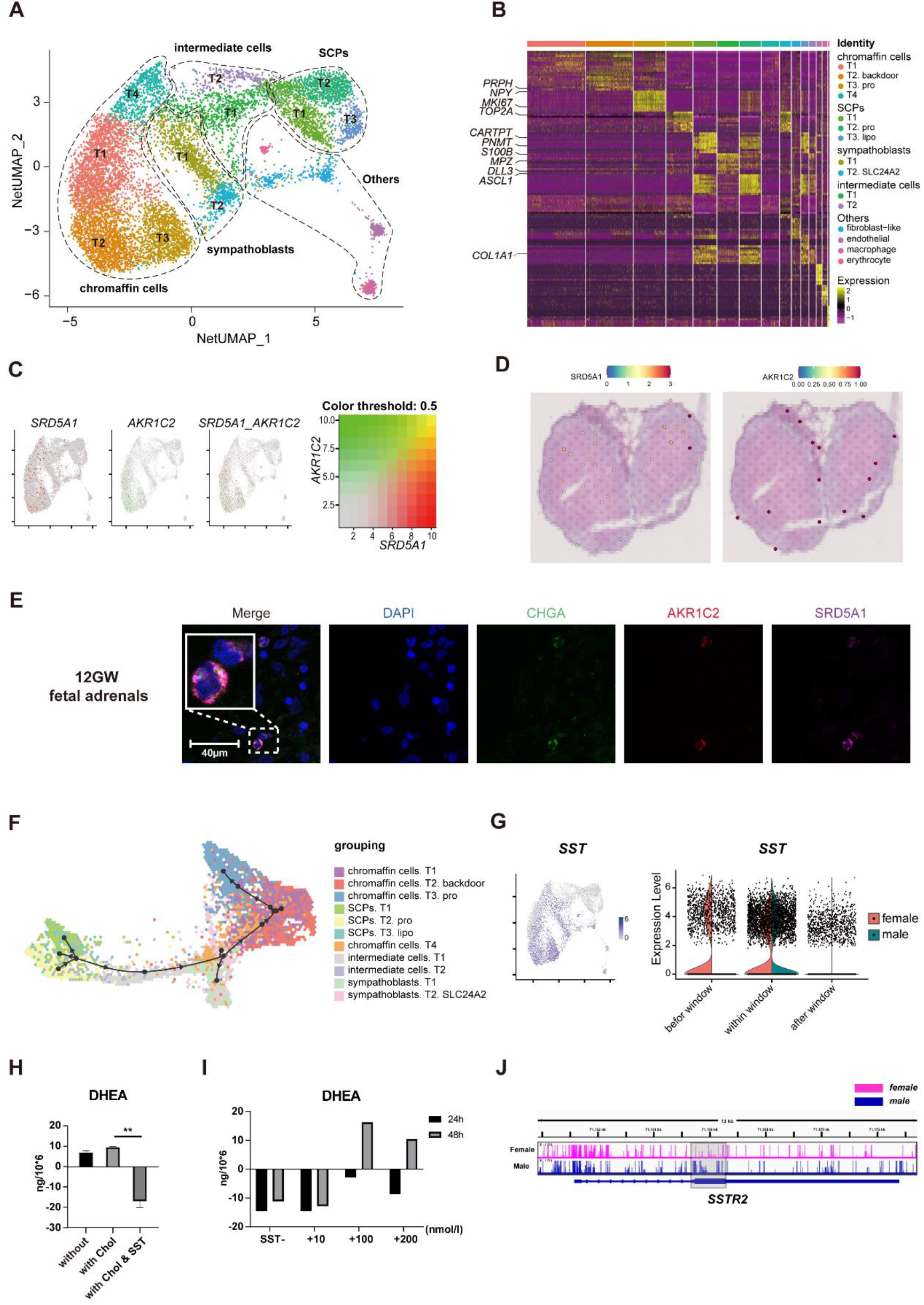
A landscape and characteristics of fetal adrenal gland neurocytes. A. Uniform manifold approximation and projection analysis visualization of adrenal neurocytes for 10× Genomics data (n = 10,812). B. Heat map of the top 10 differentially expressed genes (DEGs) between adrenal neurocyte populations. Detailed cell information and DEGs can be found in Table EV4. C. Expression patterns of *SRD5A1* and *AKR1C2* exhibited by feature plot visualization, which are key enzymes of the DHT “backdoor pathway”. A gradient of gray, red, or green indicates low to high expression, and yellow indicates coexpression. D. Visualization of the spatial transcriptome shows the locations which high express *SRD5A1* and *AKR1C2* in 8GW fetal adrenals. E. Immunofluorescence staining of SRD5A1 (purple), AKR1C2 (red), and CHGA (green) in fetal adrenal tissues. Scale bar, 20 μm. F. Differentiation of trajectory of professional steroidogenic cells using Dyno. Arrow direction indicates the trajectory of cell differentiation. G. Feature plot visualization of *SST* in fetal adrenal neurocyte data, mainly expression in mature neurocytes (chromaffin cells and sympathoblasts). Violin plot of *SST* expression with gender differences. H. ELISA of DHEA levels in the supernatant of the *in vitro* cultured fetal adrenal cells with or without somatostatin (100 nmol/l) and cholesterol (0.1 mmol/l) intervention. Chol, cholesterol; SST, somatostatin. I. ELISA of DHEA in the supernatant of the *in vitro* cultured fetal adrenal cells with different concentrations of somatostatin (−, +10 nmol/l, +100 nmol/l, +200 nmol/l). J. The landscape of chromatin accessibility site in SSTR2, of human fetal adrenal MC2R+ steroidogenic cells (female, purple; male, bule).

The chromaffin cells expressed the marker genes *CHGA* and *PHOX2B*. In addition, the T2 chromaffin cells specifically expressed *SRD5A1* and *AKR1C2*, and these cells may synthesize the active androgen dihydrotestosterone (DHT) through a “backdoor pathway” (Fig 4A-C)(Miller & Auchus, 2019, Reisch, Taylor et al., 2019). there expression levels were confirmed by immunofluorescence staining (Fig 4E).

T2 SCPs expressed the neural progenitor marker genes *S100B* and *SOX10*, the myelination-associated gene *MPZ*, and the cell cycle-associated genes *MKI67* and *TOP2A*, indicating that these cells are active proliferative cells (Fig 4B and EV2A; Table EV4). Trajectory analysis revealed that the chromaffin cells and sympathoblasts in humans originated from a cluster of SCPs(Dong et al., 2020) (Fig 4F and EV2B). This finding is highly consistent with that of previous research in mice(Furlan et al., 2017), and two intermediate cell subtypes were found to serve as a continuous “bridge” between the SCPs and mature neurocytes (Fig EV2B). These “bridge” cells expressed the SCP marker gene *HOXB9*, the functional neurocyte genes *CHGA* and *PHOX2B*, and the transient genes *ASCL1* and *DLL3* (Fig 4B and EV2B; Table EV4). Along with neurocyte development and maturation, the cell motility gene *ACTC1* and the neurodevelopmental genes *ASCL1*, *DLL3*, *GPC3*, and *HOXB9* (Furlan et al., 2017) were downregulated, whereas the cell mitosis-related genes *HSPA1B* and *PROX1* and the insulin mediator *IRS2* were upregulated (Fig EV2C).

In exploring the steroid associated regulator, we found that *SST* was highly expressed in both male and female mature neurocytes within the window of sexual differentiation and was also highly expressed before the window in females (Fig 4G) while the receptor gene *SSTR2* was higher in male steroidogenic cells (Fig 4J) Immunofluorescence staining showed that the numbers of *SST*+ neurocytes were relatively high during the window of sexual differentiation (Fig EV2D-F). *SST* encodes somatostatin, which inhibits the adrenal steroidogenic cell synthesis of DHEA, as confirmed by *in vitro* adrenocortical organoids culture test (Fig 4H). Somatostatin inhibits DHEA synthesis in primary fetal adrenal cells at low concentrations and promotes DHEA synthesis at high concentrations due to the inhibition of feminization (Fig 4I). The spatiotemporal expression of *SST* and the sex differences of *SST* reactions in sexual differentiation warrant further research.

### CD5L+ Macrophages Participate in Steroidogenesis

The immune cells were further divided into 16 distinct subtypes (Fig EV3A-B; Table EV5), which included 5 macrophage subtypes (*CD68*+ *CD163*+), 2 NK/T cell subtypes (*NKG7*+ *CD3D*+), proliferative cell subtypes (*MKI67*+ *TOP2A*+), and yolk sac-derived myeloid-biased progenitors (YSMPs, defined by *CD34*+ *MPO*+ *AZU1*+)(Bian et al., 2020). Various subtypes of immune cells were found to express lipid regulatory factors such as *APOE* and *LIPA*, and steroid biosynthesis factors, such as *POMC*, *IGF1*, and *NR4A3*. (Fig EV3C). Macrophages were the most abundant immune cells, and T4 macrophages especially expressed *CD5L* (Fig EV3A-C). T4 cells were found to be involved in cellular lipid catabolic processes and regulation of plasma lipoprotein particle levels for lipid metabolism (Fig 3D). *CD5L* were co-staining with the macrophage marker *CD68* (Fig EV3E). For fetal development, *ACY3*, *CX3CR1*, and *HMGA2*, which are associated with chemokines and cell migration, were downregulated, whereas genes such as *RHOB*, which are involved in mediating apoptosis in immune cells, were upregulated (Fig 3F). Notably, the steroid regulator *NR4A3* was highly expressed during the sexual differentiation window (Fig 3F) relative to the surge in DHEA secretion by the *in vitro* primary fetal adrenal cell co-cultured with macrophages (Fig 3G and H).

Macrophages were found to negatively regulate adrenal proliferation and differentiation via the TGFb signaling pathway network(Riopel, Branchaud et al., 1989) (Fig EV4H-J). Spatial transcriptome revealed that macrophages marker by *CD68* were mainly located in centroids serving as the primary site of FZ (Fig EV4I-J). In addition, the steroidogenic regulation genes which mentioned above such as *CD5L*, *LIPA*, *APOE*, *NR4A3*, *IGFBP1* are also highly expressed in this region (Fig EV3I). These cells may be involved in the phagocytosis of apoptotic steroid cells at the end of centrality migration(Ishimoto & Jaffe, 2011). Our data revealed the tissue specificity of fetal adrenal immune cells for steroidogenic regulation.

### Regulatory Network of Cell–Cell Interactions in Fetal Adrenal Glands

The microenvironment of steroidogenic function depends on the co-construction of various cells in fetal adrenal glands (Fig EV4A). CellChat analysis shows that the melanocortin signaling pathway acted as an autocrine pathway through the *POMC*– *MC2R* ligand receptor (Fig EV4B), as verified by Spatial transcriptome and immunofluorescence staining (Fig EV4C-D). NPY signaling pathway communication, especially between neurocytes and steroidogenic cells, is associated with cholesterol metabolism(Chen, Zhou et al., 2020) (Fig EV2G-H). The AGT signaling pathway is found between vascular myocytes and steroidogenic cells, suggesting a specific effect on sterol blood pressure(Lumbers & Pringle, 2014) and sterol insulin-resistant responsive cells(Pahlavani, Kalupahana et al., 2017) (Fig EV4E-F). The AGT-AGTR1 interaction can be demonstrated in Spatial transcriptome (Fig EV4G).

### Gonadal Non-reproductive Cells Exhibit Diverse Functions

To investigate the role of adrenal steroids in gonadal sex hormone synthesis for guiding sexual differentiation, we obtained 17 cell groups from the 10× Genomics gonads data (Fig 5A-C; Table EV6). The groups were subsequently annotated using known gene markers, such as *FOXL2* for granulosa cells, *AMH* for Sertoli cells, *INSL3* for Leydig cells, and *ARX* for somatic progenitor cells(Rotgers et al., 2018) (Fig 5B).

**Figure 5.**
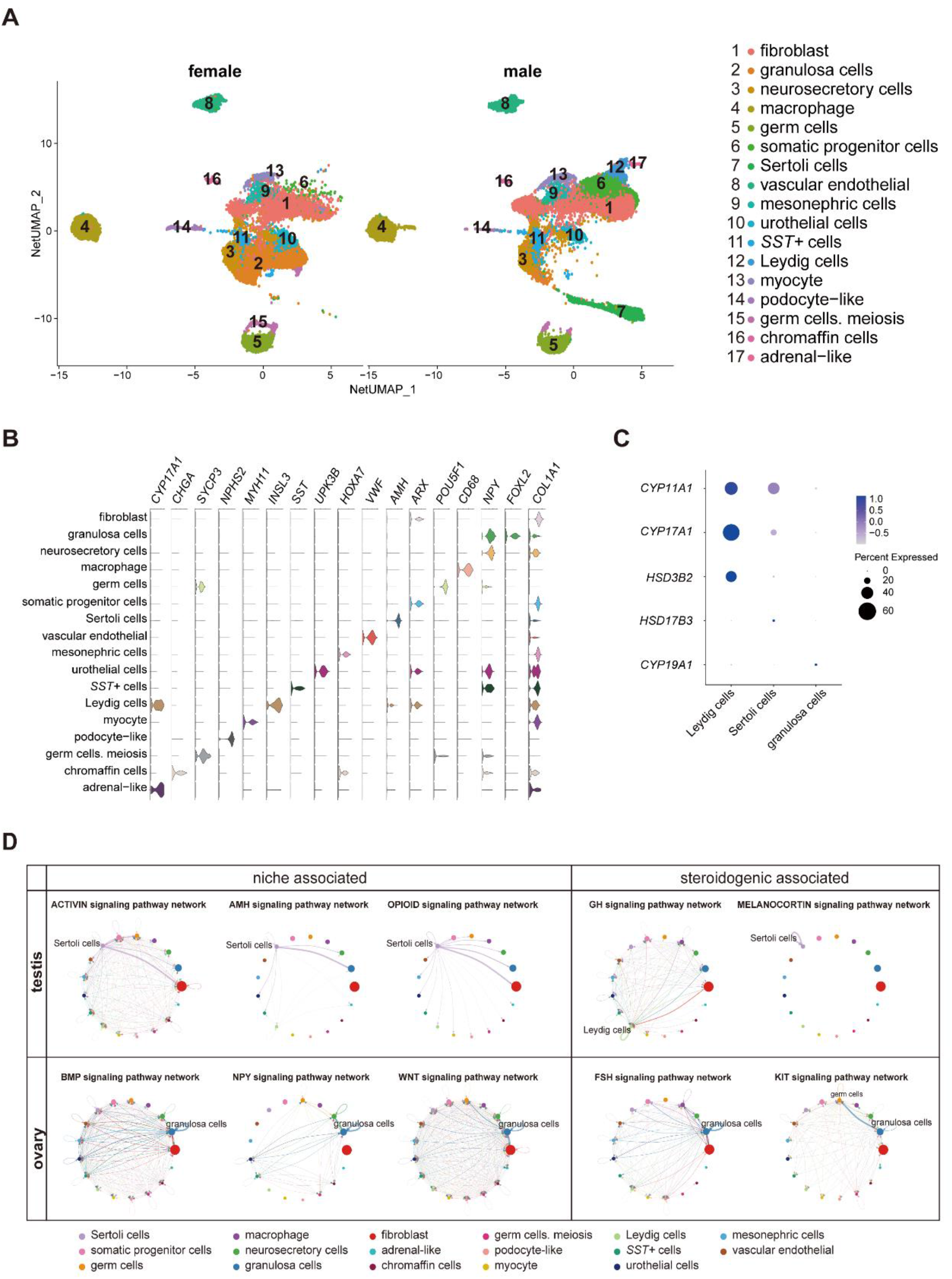
Transcriptomic landscape of human fetal gonads. A. Uniform manifold approximation and projection analysis of the transcriptomes of all-stage fetal gonadal cells split by gender (n = 53,508). B. Violin plot overview of the expression of selected marker genes by gonad clusters. Detailed cell information and differentially expressed genes can be found in Table EV6. C. Dot plot showing the expression of key enzymes for sex hormone biosynthesis (*CYP11A1*, *CYP17A1*, *HSD3B2*, *HSD17B3*, and *CYP19A1*) in different gonadal somatic cells. D. CellChat analysis showing the cell–cell interaction of gonadal somatic cells in the testis (up). The signal interaction pathway networks are divided into two functions: niche-associated (activin, AMH, opioid) and steroidogenic-associated (GH, melanocortin). The cell–cell interaction of gonadal somatic cells in the ovary (below). The signal interaction pathway networks are divided into two functions: niche-associated (BMP, NPY, WNT) and steroidogenic-associated (FSH, KIT).

The cell interactions in gonadal somatic cells (Sertoli cells, Leydig cells, and granulosa cells) were explored by CellChat (Fig 5D). Sertoli cells can be mediated by the activin, AMH, and opioid signaling pathways to form testicular cords and provide a favorable environment for germ cell survival(Li et al., 2017, Meroni, Galardo et al., 2019). In addition, Sertoli cells can be regulated by the autocrine melanocortin signaling pathway network, which may stimulate and promote the response of Sertoli cells to sex steroid biosynthesis(Aluru & Vijayan, 2008) (Fig 5D). Leydig cells were found to be regulated by GH signaling pathways to induce steroidogenesis (Fig 5D)(Colon, Svechnikov et al., 2005). However, the GH signaling pathways can cause feminization to some extent(Banerjee, Das et al., 2019). The granulosa cell interaction signaling pathway is mainly involved in oogenesis and folliculogenesis in fetal ovaries(Li et al., 2017, Zhang, Yan et al., 2018). The WNT signaling pathway promotes granulosa cell differentiation and inhibits apoptosis; the KIT and BMP signaling pathways interact with germ cells to promote oogenesis and folliculogenesis and protect preantral follicles from apoptosis(Rotgers et al., 2018, Zhang et al., 2018) (Fig 5D). As a result of immature ovarian granulosa cell development in this period, the function of ovarian steroidogenic regulation by the KIT and FSH signaling pathways may not be reflected (Fig 5C and D).

### Sex Difference in *SST*+ Cells Indicated that the Dose-dependent Effect of Somatostatin Plays a Role in Sexual Differentiation

Interestingly, *SST* in gonads was specifically expressed in a cluster of cells (Fig 5A and B and Fig EV5A). *SST* can regulate gonadotropins and thus affect the levels of sex steroids in gonads(Nakamura, Otsuka et al., 2013). The number of *SST*+ cells in males was approximately twice as much as that in females during the window of sexual differentiation (Fig EV5B). They also showed sexual differences, similar to that previously mentioned in mature adrenal neurocytes (Fig 4G). These sexual disparities were confirmed by FACS and immunofluorescence staining (Fig EV5C–D). GO analysis identified gland development, reproductive structure development, reproductive system development, and mesenchyme development (Fig EV5E). The CellChat analysis was indicated that the SST+ cells were contact with Sertoli cells by desmosomes (Fig EV5F), which confirmed by immunofluorescence staining showed that those two types of cells were adhere closely with DSC2 (Fig EV5G). Furthermore, the peak numbers of *SST*+ cells coincided with the time of emergence of Leydig cells, thereby possibly inhibiting GH to prevent feminization(Adams, Otero-Corchon et al., 2015, Banerjee et al., 2019) (Fig 5C, D and EV5B-D). As mentioned above, somatostatin exerts dose-dependent effects on sex steroids (Fig 4I), and measurement of its levels in gonads revealed high doses in testis and low doses in ovary (Fig EV5B-D). A high dose determines the “anti-female” mechanism of somatostatin that promotes androgen biosynthesis in testis, in contrast to the ovary.

To determine the potential steroidogenesis network of human fetal adrenal glands and gonads, we summarized the results of this research: adrenal glands begin to express steroidogenic-related enzyme genes at approximately 7 GW, much earlier than testis, and testis expresses steroid patterns that support de novo or partial testosterone synthesis. An *HSD3B2* expression peak was observed in females at 10 GW. Adrenal glands may synthesize small amounts of DHT. Nevertheless, the expression of steroidogenic enzymes in ovaries is limited until 14 weeks (Fig EV6).

## Discussion

Temporarily elevated *HSD3B2* was thought to consume pregnenolone by synthesizing cortisol and then reducing DHEA output, thereby avoiding the masculinization of women(Goto et al., 2006, Melau et al., 2019). In our data, HSD3B2 was high express in female while *CYP21A2* was limit during the window of sex different, high *CYP21A2* expression was not observed even until 14 GW. On account of *CYP21A2* is one of the key enzyme of cortisol synthesis (Goto et al., 2006, Melau et al., 2019), our data didn’t support previous guesses. Our hypothesis is that the steroid end-product of *HSD3B2* is probably 11-Oxygenated androgens and not cortisol(Davio et al., 2020, Pretorius et al., 2017, Turcu et al., 2020). The few reported that the role of 11-Oxygenated androgens play in the maternal - fetal cycle in placenta. Whether the 11-Oxygenated androgens participated in sexual differentiation mediated by fetal adrenal was remain unknown(O’Shaughnessy, Antignac et al., 2019). The possibility that 11-Oxygenated androgens may prevent female fetus masculinization cannot be excluded. Further research is warranted to clarify the underlying molecular mechanisms.

A study in mice reported that the cellular origin from Schwann cell precursors is the stem cell of adrenal medulla cells, which in turn could develop into chromaffin cells or sympathoblasts. A group of intermediate cell features that expresses *Htr3a*+ and *Ascl1*+, which are transient intermediate cells, is called as “bridge cells” (Furlan et al., 2017). Notably, in the present study, the trajectory of SCP was highly consistent with that of mouse. In the trajectory analysis map is a group of *ASCL1*+ intermediate cells that act as a bridge between SCP and different chromaffin cells and sympathoblasts. Our research suggested that, although the adrenal glands between humans and mice are remarkably different, they can still be reliably studied neurocyte in mouse models.

Notably, a cluster of *SRD5A1+* and *AKR1C2+* chromaffin cells was found in fetal adrenal glands. These cells may synthesize active DHT by converting An (androsterone), a process called the androgen synthesis “backdoor” pathway, which corresponds to the conversion of testosterone to DHT via the “classic” pathway(Miller & Auchus, 2019, O’Shaughnessy et al., 2019, Reisch et al., 2019). DHT is considered to be a highly bioactive androgen and reportedly promotes the development and maturation of the nervous system(Starka, Duskova et al., 2015, Venkatesh & Monje, 2017). Thus, we surmise that DHT synthesis may promote the maturation of adrenal neurocytes.

Surprisingly, *SST* expression was observed higher in male not only in adrenal glands but also in gonads during the window of sexual differentiation. *SST*-encoding somatostatin is a GH inhibitor and is reportedly associated with gonadotropin and the development of femininity(Adams et al., 2015, Nakamura et al., 2013). The chromatin accessibility of receptor genes *SSTR2* was more highly in steroidogenic cells of males. Thus, we speculate that *SST* expression is probably involved in steroid synthesis. Test results of fetal adrenal organoids suggested that somatostatin inhibits DHEA synthesis at low concentrations (10 nmol/L). However, this inhibitory effect is reversed at high concentrations (>100 nmol/L). Hence, we hypothesize that this dose-dependent reactivity of somatostatin causes different outcomes between males and females. Females would be in a long-term inhibition, whereas the opposite is true for males due to the increase in somatostatin dose with the suppression of feminized transcription factors that ultimately promotes androgen synthesis in adrenal glands and gonads. Although this effect has been noted previously(Adams et al., 2015, Banerjee et al., 2019, Nakamura et al., 2013), it has not been observed in other adrenal or gonad single-cell sequencing studies. Further investigations, such as studies of mice with Sst conditional knockout, are needed to confirm the underlying molecular mechanism.

A group of *CD5L*+ macrophages were found in fetal adrenal glands, which were reportedly mediates lipid biosynthesis in testis(Wang, Yosef et al., 2015). Primary adrenal cell culture test showed a sharp increase in DHEA synthesis in a co-culture with macrophages. This result indicated that macrophages are involved in steroidogenic regulation and synthesis in adrenal glands.

Our study of steroidogenesis in human fetal adrenal glands and gonads during sexual differentiation at a single-cell solution not only confirmed past results but also identified novel cell populations. To sum up, these findings can be linked together to form spatiotemporal regulation network, highlighting spatiotemporal differences in the steroidogenic regulatory network in adrenal glands and gonads, which we propose as a “signal and fuel” phenomenon. *SRY* gene expression is the signal of male differentiation, whereas DHEA synthesis in adrenal glands is the fuel that promotes male fetal differentiation on the right track. Our analyses provide novel insights into cell crosstalk during sexual differentiation in human fetal adrenal glands and gonads. This study will deepen our understanding of the complex regulatory mechanism of human fetal sexual differentiation.

## Materials and Methods

### Human Samples and Quality Control

Human fetal adrenal and gonad tissues were isolated from aborted fetuses following elective surgical termination of pregnancy at the Second Affiliated Hospital of Guangxi Medical University, China. Every donor for the study provided informed consent. Gestational age was validated by ultrasound for crown rump length and fetal limb length to obtain more precise information about the age of the fetuses as described previously(Napolitano, Dhami et al., 2014). The sex of the fetus can be determined by illustrations of external genitalia or PCR *SRY* gene detection. The study was approved by the ethics committee of the Second Affiliated Hospital of Guangxi Medical University (KY-0096).

### Preparation of Cell Suspension

Fresh samples of human fetal adrenal glands or gonads were placed in RPMI 1640 medium (C11875500BT, Gibco), which contained 5% fetal bovine serum (FBS; SH30070.03, HyClone) and 1% penicillin-streptomycin (15140-122, Gibco), and quickly transported to the laboratory on ice. Then, the fetal adrenal glands and gonads were separated under a stereomicroscope (Nikon)and washed with cold D-PBS (311-425-CL, Wisent) and sliced into approximately 1–2 mm^3^ pieces. The tissues were transferred to digestion solution [0.1 mg/ml Liberase TL (5401020001, Roche) and 1 mg/ml DNase I (10104159001, Roche) in RPMI 1640], at 37°C with gentle shaking throughout (adrenal glands for 20 min and gonads for 30 min), filtered through a 100 μm cell strainer (352360, Falcon), centrifuged and resuspended in 5 ml of 1X red blood cell (RBC) lysis buffer (420301, BioLegend) for 5 min on ice. Then washed twice with D-PBS containing 1% FBS, filtered through a 40 μm cell strainer (352340, Falcon), centrifuged and resuspended in DPBS with 1% FBS. The cell number and viability were assessed by Trypan blue (152 50-061, Gibco) staining and counting in a counting chamber (717805, Brand).

### Single-cell RNA-seq Library Preparation and Sequencing

We followed the approach of our previous studies(Liao, Yu et al., 2020, Yu, Liao et al., 2019). Reverse transcription and library preparation were performed using the 10x Genomics Single Cell v3 kit following the 10x Genomics protocol. Briefly, we added the single-cell suspension, gel beads and partitioning oil to 10x Genomics Chromium chip B and ran the Chromium Controller. After water-in-oil generation, samples were transferred into a PCR tube, and reverse transcription was performed using a T100 Thermal Cycler (Bio-Rad). Then, cDNA purification and library preparation were performed as the user guide. cDNA libraries were sent to Genergy Biotech (Shanghai) and sequenced by NovaSeq 6000 (Illumina).

### Spatial transcriptome library preparation and sequencing

Fetal adrenal gland tissue samples were embedded in optimal cutting temperature compound and stored at −80 °C in a sealed container. The RNA Integrity Number of fetal adrenal gland tissue collected tissue sections should be ≥ 7 before placing the tissue sections on Visium Spatial slides. A temperature of cryostat setting of –20°C for blade and –10°C for the specimen head was recommended. Tissue blocks were cut into 10μm sections and processed the capture areas using the Visium Spatial Gene Expression Kit (10x Genomics) according to the manufacturer’s instructions. Fetal adrenal gland tissue permeabilization condition was optimized using the Visium Spatial Tissue Optimization Kit. Sections were stained with H&E and imaged using Nikon Eclipse Ti2 microscop magnification, then processed for spatial transcriptomics by consulting the Visium Spatial Gene Reagent Guidelines Technical Note (CG000239) for more information. After the library construction, Agilent 2100/LabChip GX Touch was used to detect the fragment length distribution of the library. Also, q-PCR technique was used to accurately quantify the effective concentration of the library. The effective concentration of the library aimed > 10nmol/L. The qualified library was sequenced on an Illumina Novaseq 6000 platform. Cycling condition(Sequencing base length) were set 28, 90 and 10 for read1, read2 and read 3 (i7 index), respectively.

### COOL-seq library preparation and sequencing

The library preparation of COOL-seq was according to previous reports by Tang’s lab(Guo et al., 2017). After in vitro methylation of the bulk *MC2R*+ cells nuclei with M.CviPI as previous Tang’s protocol(Guo et al., 2017), the genomic DNA released by proteinase treatment was used to construct the COOL-seq library using a PBAT strategy, as previously described(Guo, Yan et al., 2015, Smallwood, Lee et al., 2014). Briefly, the bulk cells genomic DNA was bisulfite converted using the MethylCode Bisulfite Conversion Kit (Invitrogen) following the manufacturer’s instructions. Then, the purified DNAs were annealed using random nonamer primers with a 5′-biotin tag (5′-Biotin-CTACACGACGCTCTTCCGATCTNNNNNNNNN-3′) in the presence of Klenow fragments (3′-5′ exo-, New England Biolabs). Then, the primers were digested by exonuclease I (NEB) and the DNA was purified using Agencourt Ampure XP beads (Beckman Coulter). Dynabeads M280 (Invitrogen, streptavidin-coupled) were then used to immobilize the newly synthesized biotin-tagged DNA strands, and the original bisulfite-converted DNA templates were removed. Second DNA strands were synthesized using Klenow fragments with random nonamer primers (5′- AGACGTGTGCTCTTCCGATCTNNNNNNNNN-3′). After washing, the beads were used to amplify libraries using 13 cycles of PCR with the Illumina Forward PE1.0 primer and Illumina Reverse indexed primer (New England Biolabs) in the presence of Kapa HiFi HS DNA Polymerase (Kapa Biosystems). The amplified libraries were purified with Agencourt Ampure XP beads twice and were assessed on the Fragment Analyzer (Advanced Analytical Technologies). Finally, libraries were pooled (quantified with qPCR) and sequenced on the Illumina HiSeq 2500 sequencer for 150 bp paired-end sequencing.

### PCR

DNA was extracted from tissue samples according to the kit instructions (DN10, Aidlab). Fragments of the exons of *SRY* genes were amplified by CWBIO 2*ES Tap Master Mix (Dye) (CW0690M, CWBIO) with primers (forward 5′- CAGGATAGAGTGAAGCGACC-3′ and reverse 5′- CATAAGAAAGTGAGGGCTGTAAG-3′) in a 50 μl reaction mixture. PCR was performed using the Bio-Rad T100 Thermal Cycler by PCR program: 94.0°C for 2 min; 50 cycles (94.0°C, 30 sec; 60.0°C, 30 sec; 72.0°C, 30 sec), and 72.0°C for 2 min.

### Immunofluorescence Staining

The following protocols were used for staining of the fetal adrenal glands or gonads: paraffin-embedded, 4-µm formalin-fixed tissue sections were dewaxed in xylene and rehydrated with distilled water. The sections were treated with EDTA at pH 8.5 (C1034, Solarbio) to induce epitope retrieval by heating. After the samples were blocked for 30 min in phosphate buffered saline (PBS, SH30256.01, HyClone) with 2% bovine serum albumin (BSA, 9048-46-8, Sigma), sections were incubated overnight at 4°C with primary antibodies. The sections were washed three times with PBS for 5 min each and then incubated with the second antibody. for 1 h at room temperature. The detailed antibody information can be found in Table EV7. The slides were washed three times in PBS for 5 min each, and then, the nuclei were stained with DAPI (4083S, CST). Fluorescence images were captured using a laser scanning confocal microscope (TCS SP8, Laika Microscope System Shanghai Trading Co., Ltd.) and then processed with ImageJ software (NIH).

### Flow Cytometry Analysis and Cell Sorting

The cells were suspended in PBS (containing 1% BSA), then incubated with antibody (CD68, 1:100, 14-0688-82, Invitrogen or MC2R, 1:200, NB100-93419, Novus) at 4°C for 30 min. CD68 was used for sorting macrophages (Bian et al., 2020)then wash twice with PBS containing 1% BSA. Then, the cells were incubated with Alexa Fluor 594-conjugated donkey anti-mouse IgG (1:500, ab150112, Abcam) at 4°C in the dark for 30 min, then wash twice with PBS containing 1% BSA. The cells were then resuspended in PBS containing 1% BSA for FACS. All samples were loaded on a BD FACS Melody for flow cytometry cell sorting. After sorting, CD68-positive cells were used for cell coculture and MC2R-positive cells were used for multiple omics research.

Fetal gonadal cells a were fixed with fixation buffer (420801, Biolegend) at room temperature and in the dark for 20 min, then washed twice with 1X Intracellular Staining Perm Wash Buffer (421002, Biolegend). The cells were incubated with antibody (SST, 1:100, MA5-16987, Invitrogen) at 4°C for 30 minand washtwice with 1X Intracellular Staining Perm Wash Buffer. Then, the cells were incubated with Alexa Fluor 647-conjugated donkey anti-rat IgG antibodies (1:500, ab150155, Abcam) at 4°C in the dark for 30 min and wash twice and resuspended with 1X Intracellular Staining Perm Wash Buffer.. All samples were loaded on a BD C6 Plus for flow cytometry analysis.

### Cell Culture

Firstly, primary cell populations isolated from fetal adrenal glands were cultured in in 6-well plates (3516, Corning), with 2 ml of medium in RPMI 1640 with 10% FBS (SH30070.03, HyClone), 1% penicillin/streptomycin (15140-122, Gibco) and 2 mM L-glutamine (25030081, Gibco) at 37°C in a humidified 5% CO_2_ atmosphere. After 7 days, expanded adherent cells were trypsinized and reseeded in DMEM/F12 supplemented with 10% FBS, 5% horse serum (16050-122, Gibco), 100 μg/ml primocin (ant-pm-1, InvivoGen), 100 ng/ml recombinant human FGF2 (100-18C, PeproTech), and 2 mM L-glutamine(Poli, Sarchielli et al., 2019). Primary cell populations were stained with CYP17A1 and MC2R to confirm that they were steroidogenic cells.

### Cell Stimulation and Coculture

Steroidogenic fetal adrenal cells were incubated in cultivation medium in each well with or without 100 nmol/l SST (hor-299-a, ProSprc-Tany) and 0.1mmol/l cholesterol (C3045, Sigma) intervention for 24 h(Damsteegt, Hassan et al., 2019). The same cells separately were cultivated in standard cell culture medium as a control. Steroidogenic fetal adrenal cells were cultured with different SST concentrations (-, +10 nmol/l, +100 nmol/l, +200 nmol/l) of intervention. Basal DHEA and testosterone (T) were measured in the supernatants of cell culture after 24h of cultivation.

Macrophages and steroidogenic fetal adrenal cells were cocultured in 24-well Transwell chambers (3422, Corning). Steroidogenic fetal adrenal cells were incubated in the upper chamber at 6×10^4^ cells/well, and macrophages were inoculated in the lower chamber at a density of 4×10^4^ cells/well. The macrophage-free group was used as a control. Basal DHEA was measured in the supernatants of cell culture after 24 h of cultivation. The concentrations of DHEA and T in cell culture supernatants were detected by ELISAs.

### ELISA

Quantification of hormones was measured by ELISA with following kits: DHEA ELISA kit (KJ-0766, Jingsu Kejing Biological Technology) and testosterone ELISA kit (KJ-0779, Jingsu Kejing Biological Technology), according to the manufacturers’ instructions.

### Processing of Single Cell RNA-seq Data

Raw data were demultiplexed using the mkfastq application (Cell Ranger v3.1.0) to generate Fastq files. Fastq files then ran with count application (Cell Ranger v3.1.0) using default settings, which perform alignment (using STAR aligner, aligned to the GRCh38 human reference genomic data), filtering and UMI counting. UMI count tables were used for further analysis.

### Visium spatial transcriptomics data processing

Reads were demultiplexed and mapped to the reference genome GRCh38 using the Space Ranger software v.1.0.0 (10x Genomics). Count matrices were loaded into Seurat v.3.1.1 and for all subsequent data filtering, normalization, filtering, dimensional reduction and visualization. Data normalization was performed on independent tissue sections using the variance-stabilizing transformation method implemented in the SCTransform function in Seurat.

### Processing of COOL-seq data

Based on Tang’s previous research(Guo et al., 2017), raw reads were trimmed to remove the first 9 bases and to remove the adapter-contaminated and low-quality reads uing Trim Galore (v0.3.3). The cleaned and QC-ensured reads were then aligned to the in-silico bisulfite-converted human genome reference (hg19) using Bismark (v0.7.6) with paired-end and non-directional mapping parameters. After paired-end mapping, the unmapped reads were re-aligned to the same reference genome in single-end mode. Duplicated reads from the PCR amplification step were identified and removed by using their genomic coordinates under published SAMtools (v0.1.18) following the “samtools rmdup” command (v0.1.18) parameter: “samtools rmdup” for paired-end reads and “samtools rmdup -s” for single-end reads. The process about quantification of WCG and GCH methylation level were according to the previous Tang’s research(Guo et al., 2017). The visualization of methylation and chromatin accessibility were based on The Integrative Genomics Viewer (IGV, v2.5.x) software.

### Cell-type Identification and Dimensionality Reduction

The “Seurat” package (v.3.1.1)(Satija et al., 2015) was used as the first analytical package. For 10x Genomics data, UMI count tables from both replicates from all fetal adrenal gland and gonad samples were loaded into R using the ‘Read10X’ function, and ‘Seurat’ objects were built from each sample. Each object was filtered and normalized with default settings. Specifically, cells were retained only if they contained > 200 expressed genes and < 4,500 genes and had < 15% reads mapped to the mitochondrial genome. After the cell filtration of each object, we used the ‘merge’ function from “Seurat” to combine the objects into two main objects according to the source of organs: adrenal and gonad, for downstream analysis. Cells were normalized to the total UMI read counts by the ‘NormalizeData’, ‘FindVariableFeatures’ and ‘ScaleData’ functions, guided by tutorial in http://satijalab.org/seurat/. Then, the objects were explored with the view of molecular networks for removing the batch effect by the ‘RSCORE’ function from the “RSCORE” package(Dong, Zhou et al., 2019). Principal component analysis was performed by the ‘RunPCA’ function from “Seurat”, and the top 35 principal components were selected for UMAP analysis. The UMAP(Becht, McInnes et al., 2018) analysis was performed by the ‘RunUMAP’ function from “Seurat”. Similar cells were clustered and detected using the Louvain method for community detection by the ‘FindNeighbors’ function. Discrete clusters from adrenal data and gonad data were detected using ‘FindClusters’ and annotated by specific cell markers from the “cellmarker” database (http://bio-bigdata.hrbmu.edu.cn/CellMarker/). The erythrocytes (*HBB*, *ALAS2*) were filtered out by the ‘subset’ function. After removal of erythrocytes, 75,482 adrenal cells and 53,508 gonad cells were retained for downstream analysis.

The screened cells were reclustered using the same analytical parameters. Forty-two discrete clusters from adrenal data and 25 from gonad data were detected. The clusters were annotated on the basis of feature genes. Finally, the clusters were regrouped into the main cell groups by definition and similarity.

### Cell Cycle Analysis

As in our previous studies(Yu et al., 2019), cell cycle analysis was performed by the “Seurat” package by using previously defined cell cycle genes. We calculated a ‘cycle score’ for each cell based on the expression of cell cycle genes. Cells were considered aperiodic if the cycle score was less than two; otherwise, they were considered proliferating.

### Gene Ontology (GO) and KEGG Analysis

To detect differences in gene function expression among cell clusters in single-cell data, we generated GO gene sets using datasets from “Seurat” single-cell object conversion. GO enrichment analysis was conducted using the “clusterProfiler” package (V3.12.0)(Yu, Wang et al., 2012). Before analysis, we transferred gene names from ‘symbol’ to ‘entrezid’ according to the “org.Hs.eg.db” package (V3.8.2)(Carlson, 2019). Terms with a p-value < 0.05 were considered and enriched as significant. Dot plots and gene-concept networks were illustrated using the “enrichplot” package (V1.4.0)(Guangchuang Yu, 2019).

### Developmental Pseudotime Analysis

Detailed pseudotime for different cell types was performed using the “Dyno” package (v2.10.1)(Saelens, Cannoodt et al., 2019) following the guideline settings. The inference methods in Dyno were wrapped within “Docker” containers (available at https://methods.dynverse.org). First, the ‘wrap_expression’ function was used to generate Dyno objects by transforming ‘Seurat’ object counts and normalized expression data. Then, the Dyno objects were preset to a “start_id” as an initial state of the cell by the “add prior information” function. In addition, the “add_grouping” function was used to add the results of the cell definition from ‘Seurat’ before the trajectory analysis. Finally, the “PAGA tree” method(Wolf, Hamey et al., 2019) was selected for trajectory analysis based on the characteristics of our data. Visualization of trajectory analysis was shown by the ‘plot_dimred’ function. All trajectory analysis models were constructed in the correct biological context.

### Cell-Cell Interaction Network Analysis

The network analysis of signal interactions between cells was performed using the “CellChat” package(Jin, Guerrero-Juarez et al., 2021). The CellChat data were generated from the previous ‘Seurat’ object content by the ‘data.input’ function. The ‘idents’ in the CellChat data were obtained by the cell labels in Seurat objects. ‘Secreted signaling’ and ‘cell-cell contact’, two models of cell interaction, were pulled out from ‘CellChatDB’ for our follow-up analysis by the ‘CellChatDB.use’ function. In the subsequent interpretation of the “CellChat” analysis results, we considered the biological background and ignored some unreasonable results in the gonad data.

### Data availability

Processed and raw human single cells sequencing data are available via the Gene Expression Omnibus (GEO) (GEO: GSE167860).

## Acknowledgements

We thank the donors for participating in this study. We thank The Second Affiliated of Guangxi Medical University for support sample collection; H. Mi, L. Mo from The First Affiliated Hospital of Guangxi Medical University, H. Bai, M. Wei, Z. Liu, D. Peng, Y. Su, S. Yi, from Gynaecology and Obstetrics Hospital of Guangxi Zhuang Autonomous Region, Q Meng, C. Huang, Y. Xie, D. Li, P. Wei from Guangxi Medical University, X. Sun from The Third Affiliated Hospital of Guangzhou Medical University for experimental help; This work was supported by grants from the National Key Research and Development Program of China (2017YFC0908000), the Natural Key Research and Development Project (2020YFA0113200), the Major Project of Guangxi Innovation Driven (AA18118016), the Guangxi key Laboratory for Genomic and Personalized Medicine (16-380-54, 17-259-45, 19-050-22, 19-185-33, 20-065-33), and the Natural Science Foundation of China (81770759, 82060145, 31970814).

## Author Contributions

Y.W. Y.J., and Z.M. contributed to the study design. B.G., N.Q., Y.G. and Y.L. mainly performed cell suspensions, 10x single cell RNA-seq Library preparation. Y.G., N.Q. and Z.C. in PCR analysis. B.G., Y.G., Y.W. and N.Q. in immunofluorescence staining and confocal imaging. Y.G. performed cell stimulation, coculture and hormone quantification in vitro experiment. B.G. together with N.Q performed flow cytometry analysis and cell sorting. Y.W., Z.C., J.J., Y.Y., C.S. and Q.Z. collected fetal samples and patient information. Y.H, H.C, L.Z, M.L, J.C provided clinical technical guidance. Y.W., Y.Y, X.M contributed to bioinformatic analyses. Y.W and Y.J. contributed to manuscript preparation. Z.M. conceived and directed the study, obtained funding, revised the manuscript. All authors provided critical proposal and approved the final version of the manuscript.

## Conflict of interest

The authors declare that they have no conflict of interest.

---

**Figure EV1.**
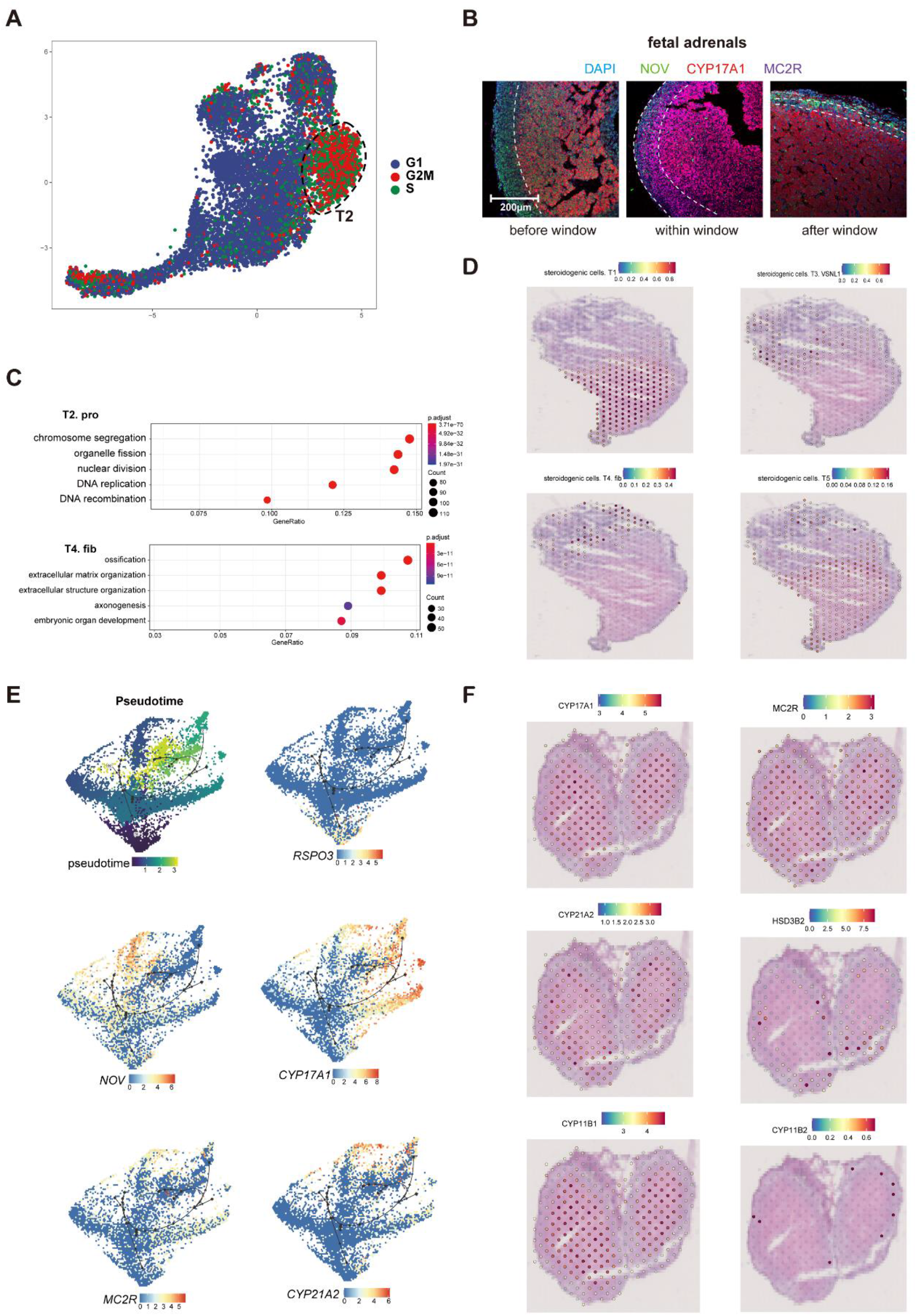
Dynamic gene expression patterns of fetal adrenal steroidogenic cells. A. Uniform manifold approximation and projection analysis visualizations of the cell cycle score of professional steroidogenic cell populations. Cell cycle states: G1 (blue), G2/M (red), S (green). B. Immunofluorescence staining of NOV (green), CYP17A1 (red), and MC2R (purple) in adrenal glands. Scale bar, 20 μm. n = 3. C. Dot plot showing the GO functional analysis of T2 and T4 steroidogenic cluster cells. P value and percent of counts as in figure. D. Visualization of the spatial transcriptome shows the location of T1, T3. *VSNL1*, T4. fib and T5 steroidogenic cells in 9GW fetal adrenal. E. Pseudotime ordering of professional steroidogenic cells using Dyno. Expression of genes associated with the adrenal marker genes *RSPO3*, *NOV*, and *MC2R* or the steroidogenic enzymes-related genes *CYP17A1* and *CYP21A2* mapped on professional steroidogenic cells. Color scale: red, high expression; blue, low expression. F. Visualization of the spatial transcriptome shows the locations which high express *CYP17A1*, *CYP21A2*, *CYP11B1*, *MC2R*, *HSD3B2* and *CYP11B2* in 8GW fetal adrenals.

**Figure EV2.**
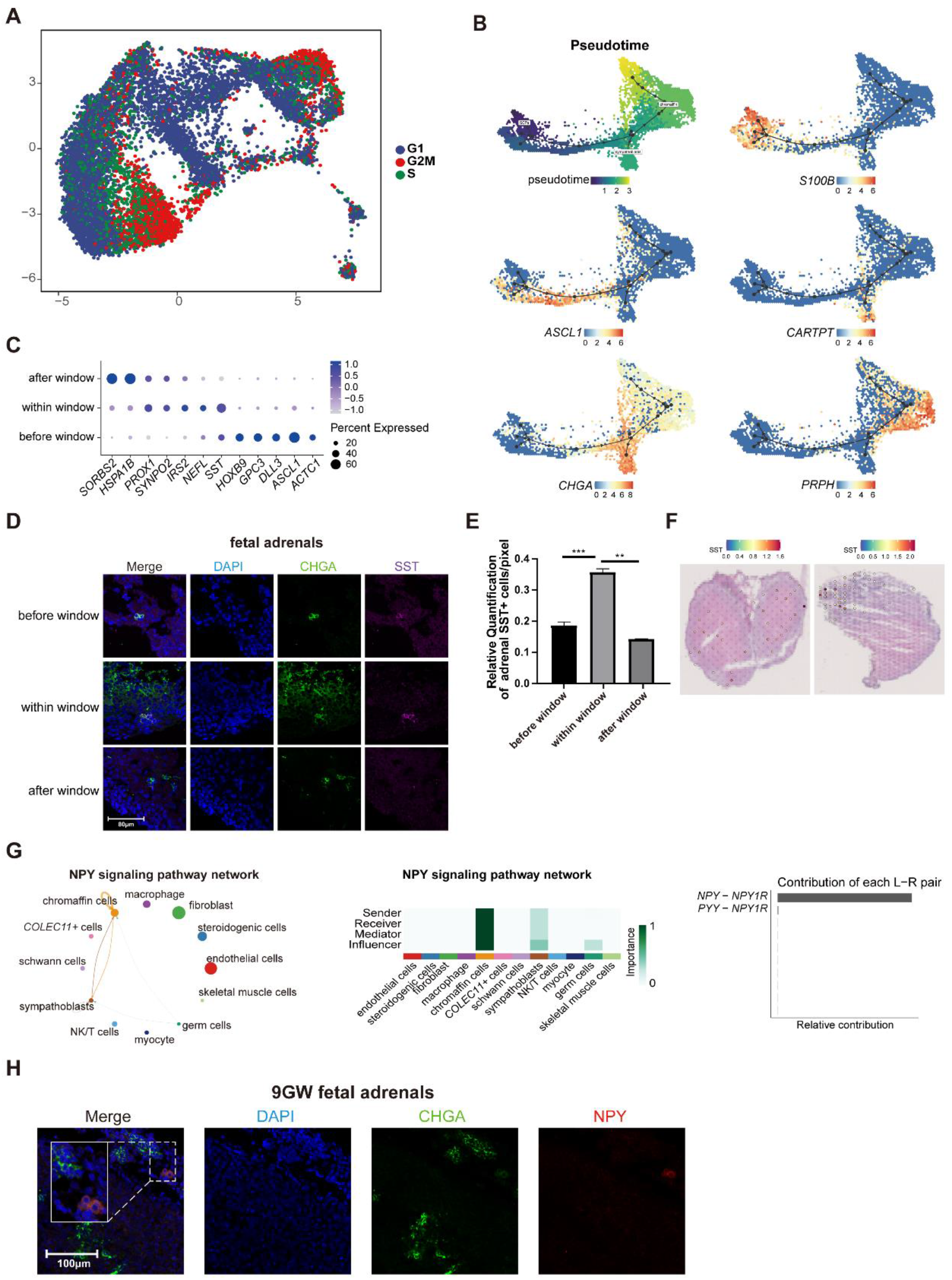
Transcriptional dynamics of human fetal adrenal neurocytes. A. Uniform manifold approximation and projection analysis visualizations of the cell cycle score of neurocyte populations. Cell cycle states: G1 (blue), G2/M (red), S (green). B. Pseudotime ordering of neurocyte populations using Dyno. Expression of genes associated with the indicated genes (*S100B*, *ASCL1*, *CARTPT*, *CHGA*, *PRPH*) mapped. Color scale: red, high expression; blue, low expression. C. Dot plots of differential gene expression of three stages of neurocyte groups (before, within, and after sexual differentiation). D. Immunofluorescence staining of SST (purple) and CHGA (green) in the adrenal glands during the window of sexual differentiation. E. Relative quantification numbers of adrenal SST+ neurocytes at different stages. Scale bar, 20 μm. n = 3. F. Visualization of the spatial transcriptome shows the locations which high express *SST* in 8GW fetal adrenals and 9GW fetal adrenal. G. CellChat analysis showing the NPY cell-secreted signaling pathway among chromaffin cells, sympathoblasts, and germ cells. Bar plot showing ligand– receptor pair contributions mainly by *NPY*-*NPY1R*. H. Immunofluorescence staining of NPY (red) and CHGA (green) in the adrenal glands. Scale bar, 20 μm.

**Figure EV3.**
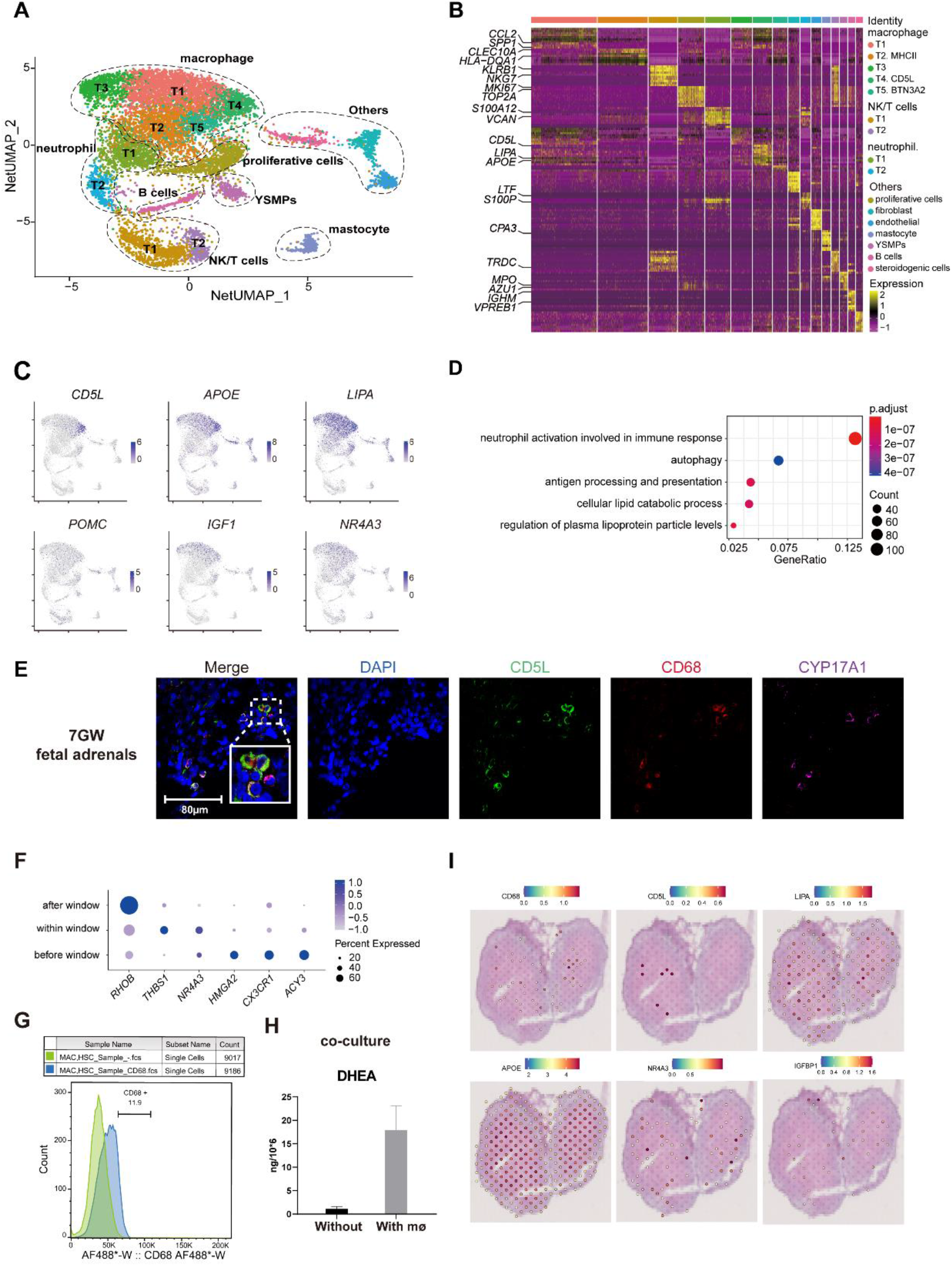
A landscape and characteristics of fetal adrenal gland immune cells. A. Uniform manifold approximation and projection analysis visualization of adrenal immune cells for 10× Genomics data (n = 10,704 cells). B. Heat map of the top 10 differentially expressed genes (DEGs) between immune cell populations. Detailed cell information and DEGs can be found in Table EV5. C. Feature plot visualization of lipid metabolic regulators (*CD5L*, *APOE*, *LIPA*, *POMC*, *IGF1*, and *NR4A3*) in fetal adrenal immune cell data. D. Dot plot displaying the GO functional analysis of *CD5L*+ macrophages. P value and percent of counts as in figure. E. Immunofluorescence staining of CD5L (green), CD68 (red), and CYP17A1 (purple) in fetal adrenal tissues. Scale bar, 20 μm. F. Dot plots of differential gene expression of three-stage immune cell groups (before, within, and after sexual differentiation). G. FACS analysis of CD68+ cells in fetal adrenal tissues at 8–12 gestational weeks (n = 4). H. ELISA of DHEA in the supernatant of the in vitro cultured fetal adrenal cells, alone or cocultured with CD68+ (macrophage)-sorted cells from fetal adrenal tissues. mø, macrophage. I. Visualization of the spatial transcriptome shows the locations which high express *CD68*, *CD5L*, *LIPA*, *APOE*, *NR4A3* and *IGFBP1* in 8GW fetal adrenals.

**Figure EV4.**
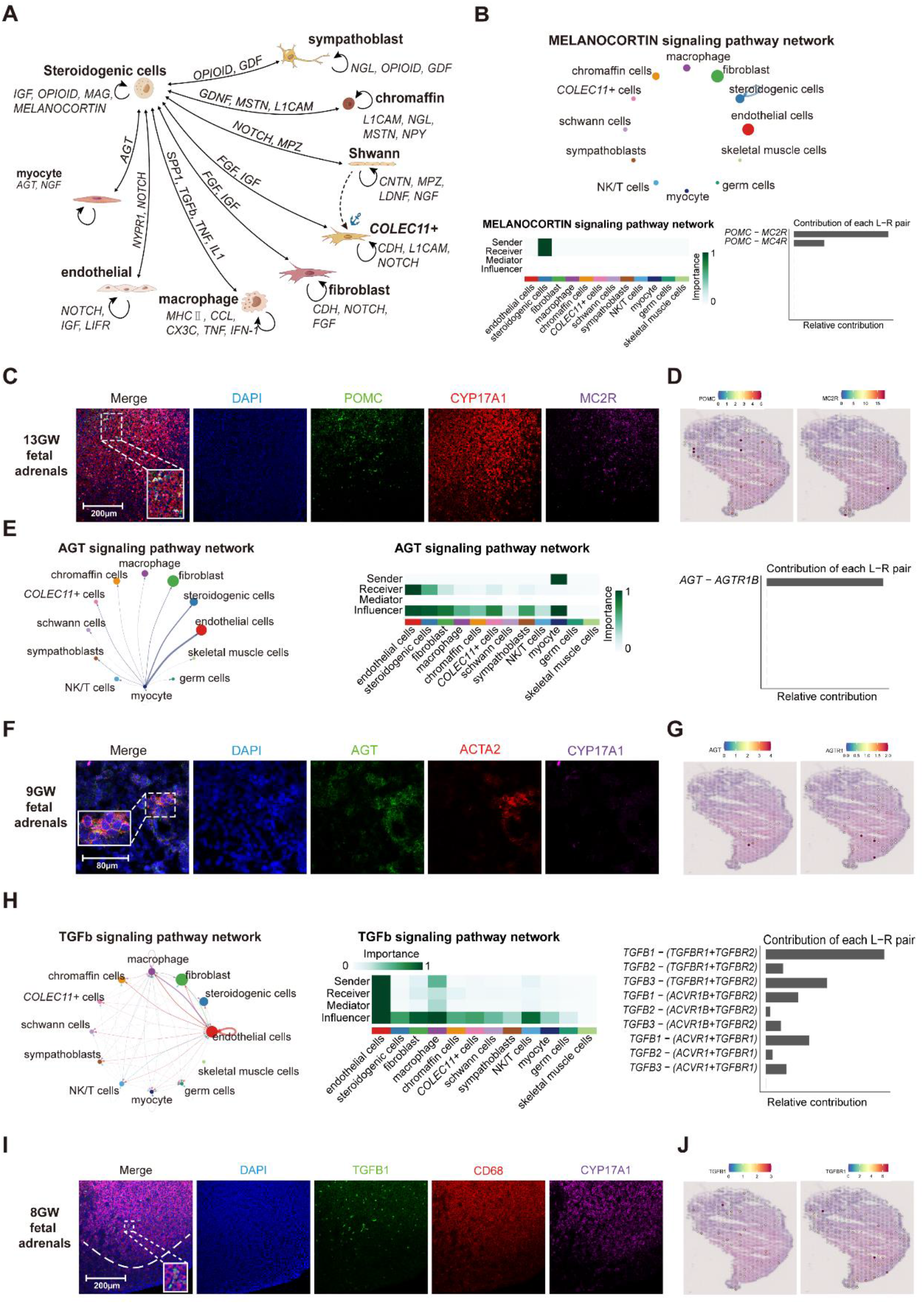
Cellular interactions in fetal adrenal glands. A. Diagram of the regulatory network between professional steroidogenic cells and other cells in the fetal adrenal glands. B. CellChat analysis showing the melanocortin-secreted signaling pathway within the professional steroidogenic cell cluster itself. Bar plot showing ligand–receptor pair contributions by *POMC*–*MC2R* and *POMC*–*MC4R*. C. Immunofluorescence staining of POMC (green), MC2R (purple), and CYP17A1 (red) in adrenal glands at 13 gestational weeks (GW). Scale bar, 20 μm. D. Visualization of the spatial transcriptome shows the locations which high express *POMC* and *MC2R* in 9GW fetal adrenal. E. CellChat analysis showing the AGT-secreted signaling pathway between myocytes and other types of cells. Bar plot showing ligand–receptor pair contribution by *AGT*–*AGTR*. F. Immunofluorescence staining of AGT (green), ACTA2 (red), and CYP17A1 (purple) in adrenal glands at 9 GW. Scale bar, 20 μm. G. Visualization of the spatial transcriptome shows the locations which high express *AGT* and *AGTR1* in 9GW fetal adrenal. H. CellChat analysis showing the growth regulator TGFb cell-secreted signaling pathway between immune cells and other types of cells. Bar plot showing ligand– receptor pair contributions mainly by TGFB1-(*TGFBR1+TGFBR2*). I. Immunofluorescence staining of TGFB1 (green), CD68 (red), and CYP17A1 (purple) in the adrenal glands. Scale bar, 20 μm. J. Visualization of the spatial transcriptome shows the locations which high express *TGFB1* and *TGFBR1* in 9GW fetal adrenal.

**Figure EV5.**
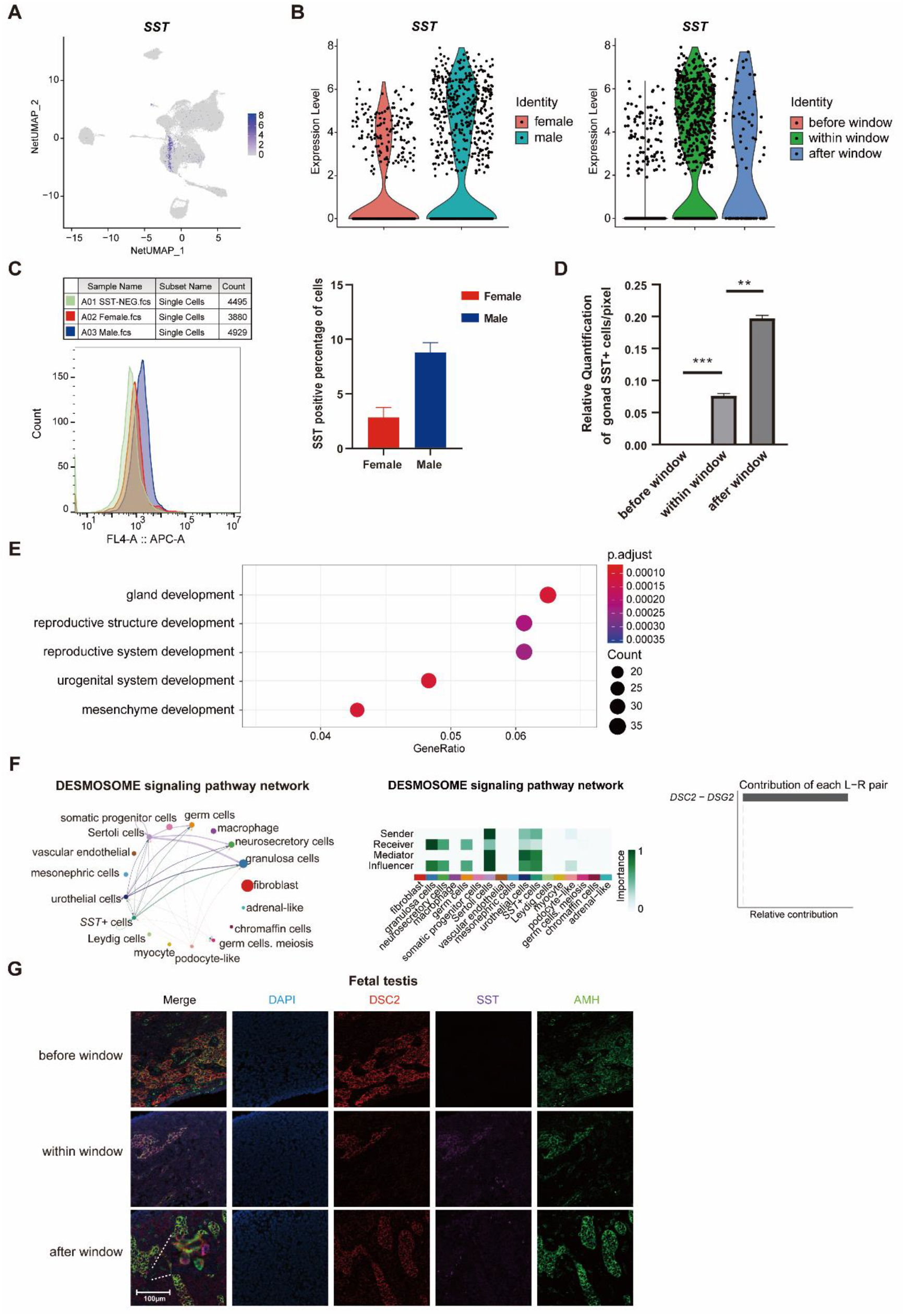
Characteristics of SST+ cells in fetal gonads. A. Featureplot showing the cluster of SST+ cells in the gonadal 10× Genomics data. B. Violin plots of *SST* gene expression patterns of female and male gonads spanning the window of sexual differentiation. C. FACS analysis of fetal gonad *SST*+ cells at 8–10 gestational weeks. Male n = 3, female n = 3. D. Relative quantification numbers of gonadal SST+ cells at different stages. E. Dot plot showing the GO function analysis of SST+ cells. P value and percent of counts as in figure. F. CellChat analysis showing the desmosome cell–cell contact signaling pathway between SST+ cells and other gonadal somatic cells. Bar plot showing ligand– receptor pair contributions by DSC2–DSG2. G. Immunofluorescence staining of AMH (green), DSC2 (red), and SST (purple) in testes spanning the window of sexual differentiation. Scale bar, 20 μm. n = 3.

**Figure EV6.**
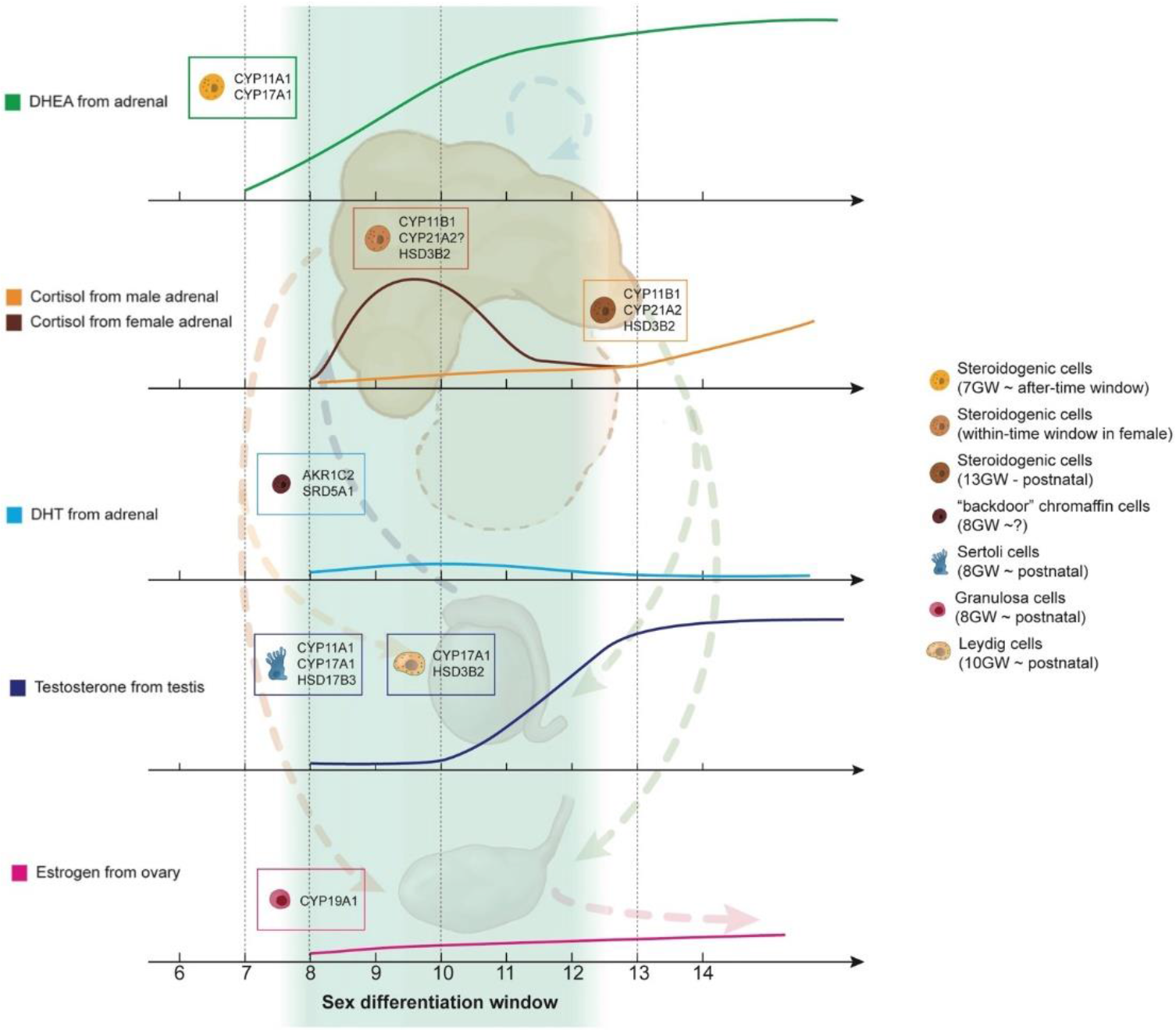
A schematic illustration showing cell-type-specific steroidogenic-related genes and steroid levels in adrenal glands and gonads.

